# How altricial birds respond to a heat challenge: organismal perspectives on coping with a future climate scenario

**DOI:** 10.1101/2024.01.11.575240

**Authors:** Mary J. Woodruff, Susanna N. Tsueda, Tiernan S. Cutrell, Ethan A. Guardado, Douglas B. Rusch, Aaron Buechlein, Kimberly A. Rosvall

**Author notes:** **Corresponding Author** Mary J. Woodruff 1001 E. 3^rd^ St., Rm A318 Bloomington, IN 47405. **Author Contributions:** Conceptualization: M.J.W. and K.A.R.; Investigation: M.J.W, S.N.T, T.S.C., E.A.G., D.B.R., and A. B.; Data curation: M.J.W., S.N.T., E.A.G., D.B.R., and A. B.; Formal Analysis: M.J.W., S.N.T., E.A.G., D.B.R., and A. B.; Project Administration: M.J.W; Supervision: K.A.R.; Writing – Original Draft M.J.W. and K.A.R. with feedback from all authors. **Data Availability:** Data will be available on the Dryad Digital Repository upon manuscript acceptance.

## Abstract

1. The ability to cope with heatwaves is likely to influence species success amidst climate change. However, relatively little is known about heat-coping mechanisms in endotherms, which are increasingly pushed to their thermoregulatory limits. We experimentally elevated nest temperatures by 4.5°C for 4 hours, focused on 12-day-old tree swallows (*Tachycineta bicolor*).
2. Nestlings exposed to sub-lethal heat moved towards cooler air at the nest box entrance, they panted more, and they weighed less than controls, suggesting panting-induced water loss. They also exhibited higher heat shock protein (HSP) gene expression in the blood, alongside widespread transcriptional differences related to antioxidant defenses, inflammation, and apoptosis. Nestlings exposed to milder heat were more likely to recruit into the breeding population, suggesting these coping mechanisms may be quite effective.
3. We also tested hypotheses on the drivers of variation in HSP gene expression, which was especially marked after heat-exposure. Even siblings in the same nest differed in HSP gene expression by over 14-fold. Heat-induced HSP levels were unrelated to individual body mass, or among-nest differences in brood size, temperature, and behavioral thermoregulation. However, nest ID explained a significant amount of HSP variation, which was larger between nests than within nests, pointing to genetic or early developmental factors
4. These results fill key knowledge gaps on thermoregulatory mechanisms in birds. We document ample individual variation upon which selection may act in the context of climate change and we underscore the need to understand intra-specific variation, an oft-ignored element that nevertheless shapes what is possible for future adaptation or acclimation to heat.

## Introduction

As heatwaves intensify (Fischer et al. 2021), researchers are mobilizing to assess which organisms are likely to persist (Moore & Schindler 2022). The potential for adaptation to climate change should relate to the scope of variation within a population; after all, heritable variation is the raw material of natural selection (Lande 1979) and individual-level plasticity shapes the pace of evolution (Fox et al. 2019).

However, studies on intra-specific, among-individual variation in heat-responses are rare, particularly in endotherms, which are increasingly pushed to their thermoregulatory limits (McKechnie & Wolf 2019).

To combat the negative effects of heat, animals adjust their physiology and behavior. Behavioral shifts may be a first line of defense (reviewed by: Huey et al. 2012; Muñoz 2022). For instance, animals may change the time of day they are active (Gilbert et al. 2022), seek cooler microhabitats (Verzuh et al.

2023), and pant or bathe to evaporatively cool (Loughran & Wolf 2020). When heat is unavoidable and body temperatures rise, animals may activate additional physiological responses, including upregulated heat shock proteins (HSPs) that minimize damage via protein refolding (Feder & Hofmann 1999; Lindquist & Craig 1988). HSPs can be co-regulated with other physiological mechanisms (Lipshutz et al. 2022), which collectively work towards restoring homeostasis (White et al. 2007). However, all of these coping mechanisms may incur costs. Behavioral thermoregulation may trade-off with foraging (Mason et al. 2017) or water balance (Albright et al. 2017), and chronically elevated HSPs may have energetic costs (Sørensen et al. 2003). Organismal perspectives that consider diverse heat responses and associated consequences are important for determining whether responses to heat are adaptive.

Responses to heat are rarely uniform within a species. Many studies have documented a*mong-population* differences in heat tolerance (Bennett et al. 2019), whereas *among-individual* differences remain poorly understood (e.g., Humanes et al. 2022), despite repeated calls to address this gap in knowledge (Huey et al. 2012; Muñoz 2022). We know, for example, that panting, basal metabolic rate, and evaporative water loss can differ dramatically among individuals (Pessato et al. 2020; Wojciechowski et al. 2021), and we need more information on the scope and drivers of such variation.

Here we begin to fill these knowledge gaps with a heat challenge that simulated a single hot afternoon we might expect within the next century of climate change (Reidmiller et al. 2018). Using 12-day-old tree swallows (*Tachycineta bicolor*) confined to their nesting cavity, our first aim was to assess the phenotypic and performance effects of punctuated heat exposure. We focused on a set of thermoregulatory mechanisms, including (a) panting, (b) space use, (c) HSP gene expression, and (d) a global analysis of other transcriptomic effects. We assessed potential consequences of coping with heat by quantifying heat effects on nestling mass, begging behavior, fledging, and recruitment into the breeding population. Our second aim was to explore nest- and individual-level traits that may contribute to within-population variation in heat tolerance. Together, these analyses assess critical parameters for understanding adaptation to climate change; we cannot expect a species to adapt to climate change, unless there is individual variation upon which selection may act.

## Methods

### Study system

Our experiment occurred in the nesting cavity – here, a human-made nest box – where nest temperatures average 12.3 ± 0.8°C above ambient (mean ± SE, control data, this study). These data mirror general observations that microhabitats can reach temperatures well above that of ambient air (Cunningham et al. 2021). Our experiment was performed in southern Indiana, USA (39.17° N, 86.53° W) from May to June 2021, when nestlings were 12 days post-hatch (D12) when nestlings are fully endothermic (Marsh 1980), have accelerating feather growth, and have typically reached asymptotic mass (McCarty 2001). Hatch day is denoted as D1, and nestlings fledge around D21 (Marsh 1980).

Average brood size was 4.5 (range: 3-6). All methods were approved by the Institutional Animal Care and Use Committee and conducted with appropriate state and federal permits.

### Experimental heating

We elevated nest temperatures using air-activated warmers (Uniheat 72hr, hereafter “packs;” *Fig. S1;* (Albert et al. 2023; Woodruff et al. 2023). Packs heat via the oxidation of iron powder when the packaging is opened and the mixture of charcoal, iron powder, vermiculite, salt, sawdust, and water is exposed to air. Because we were targeting sub-lethal temperatures, we pilot tested the use of 3-packs per nest in empty nests, finding an average elevation of 5°C (see *SI§A*). Thus, we expected nest temperatures to: (i) exceed the upper end of the expected thermoneutral zone (∼37.5°C avg for small songbirds, extracted from Appendix S1 in Wolf et al. 2017) - the point at which animals increase energy expenditure to thermoregulate (Mitchell et al. 2018), and (ii) fall below expected lethal limits that can occur with sustained exposure ≥ 45°C (Pollock et al. 2021). We heated each nest with three packs during the heat of the afternoon (start time = 11:52 ± 46 min; duration = 4.1 ± 0.2 h). In each control nest, we used three cooled packs in an identical fashion. We standardized nests and habituated birds to foreign objects, including temperature loggers, ≥ 48h in advance (see *SI§A*).

Treatments were balanced by brood size and age of the mother. Total sample sizes were n = 25 heat nests, n = 112 heat nestlings, n = 21 control nests, and n = 91 control nestlings.

We used iButton temperature loggers to assess experimental effects. One iButton was secured onto the nest cup facing down, measuring temperature at the surface of the nesting material at 10 min intervals. This iButton was along the side of the nest cup to avoid nestlings sitting directly on it. We used these data in two core analyses. (1) We assessed how quickly experimental nest temperatures increased by collating the 10 most recent pre-experiment iButton reads and the experimental temperature data. We then fit these data to a 4-parameter logistic curve, which showed temperatures dramatically rising 12 min after packs were placed in the nest box, after which temperatures plateaued (median inflection point of1.2 time bins; quartile range: 0.3-2.6; *Fig. S2*). Thus, our treatment quickly elevated and sustained temperatures for the 4h experiment. (2) We calculated mean nest and ambient temperatures across the duration of the experiment, beginning when packs were placed in the nest and continuing until nestlings were sampled. Ambient data were downloaded from the National Oceanic and Atmospheric Administration (NOAA) database, using hourly dry bulb temperature from the nearest weather station (Station ID: WBAN:03893), which was 18.5 km away at a similar elevation. NOAA data allowed us to control for ambient effects not already accounted for by counterbalancing by date. Mean ambient temperatures per nest did not statistically differ between treatments (β = 1.07, SE = 3.61, F_1,44_ = 0.77, p = 0.39; *Table 1, Fig. 1*).

**Table 1:**
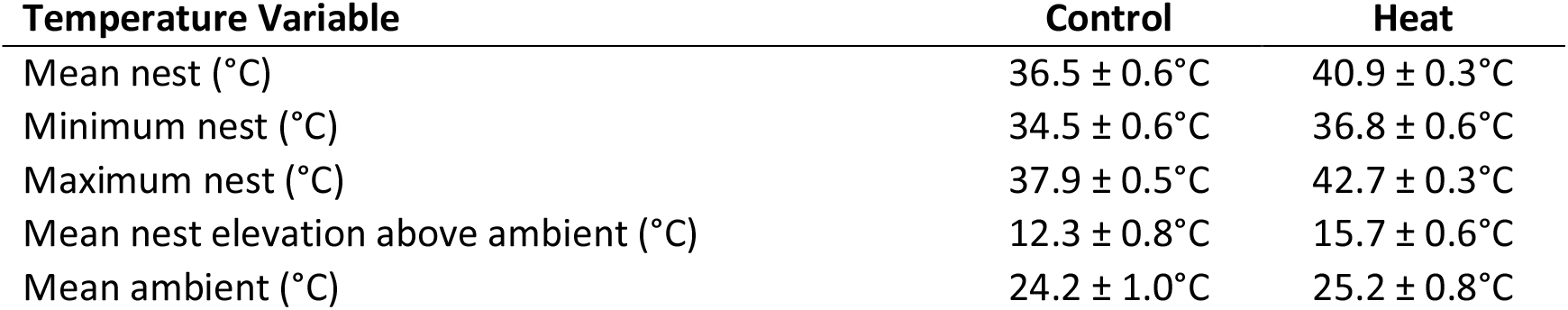
Treatment effects on temperature. Nest data come from iButton values collected at 10-min intervals, averaged per box across the experiment (± standard error). Ambient data come from NOAA values collected at 1-hour intervals, averaged per box, then by treatment.

| Temperature Variable | Control | Heat |
| --- | --- | --- |
| Mean nest (°C) | 36.5 ± 0.6°C | 40.9 ± 0.3°C |
| Minimum nest (°C) | 34.5 ± 0.6°C | 36.8 ± 0.6°C |
| Maximum nest (°C) | 37.9 ± 0.5°C | 42.7 ± 0.3°C |
| Mean nest elevation above ambient (°C) | 12.3 ± 0.8°C | 15.7 ± 0.6°C |
| Mean ambient (°C) | 24.2 ± 1.0°C | 25.2 ± 0.8°C |

**Figure 1:**
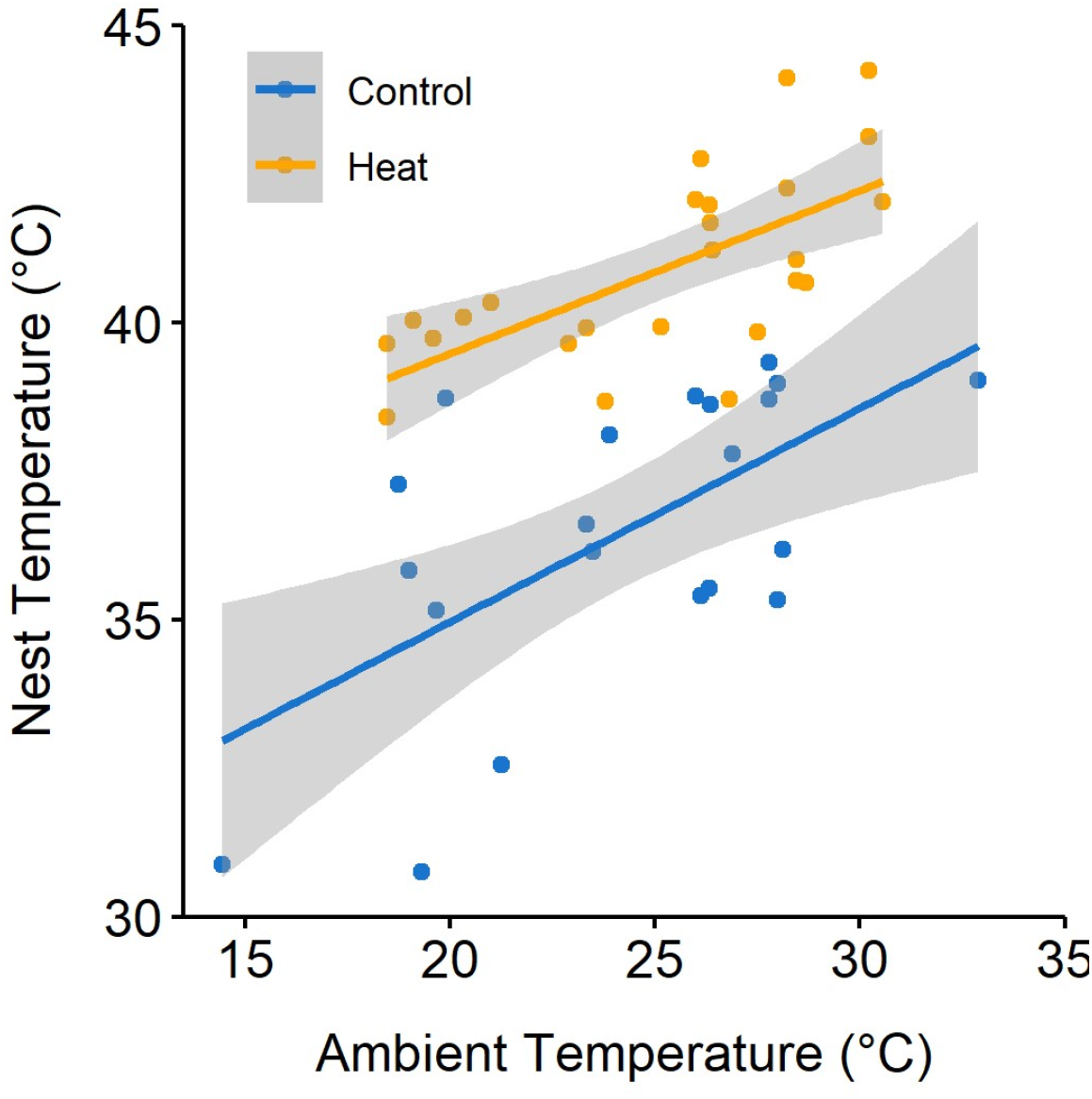
Mean nest temperature by mean ambient temperature (°C). Each point represents one nest. Shading represents 95% confidence intervals.

A second iButton was attached to the internal box wall ∼3 cm above the rim of the nest cup to measure relative humidity (RH). Mean nest RH was 54.3 ± 2.1%, and nest RH was strongly correlated with ambient RH from NOAA data (Pearson r = 0.87; *Fig. S3*). Minimum nest RH averaged 44.8 ± 2.1% and never reached below 25%, when evaporative cooling may be constrained (Van Dyk et al. 2019), so we did not consider RH further. Details in *Table S3*.

### Behavioral observations

We placed a camera (GoPro HERO Session 4) in the upper corner of the nest box at the same time as pack placement. Using JWatcher (version 1.0, Blumstein & Daniel 2007), a single observer (S.N.T.) scored behaviors from the second hour of the experiment, allowing packs to warm and birds to recover from our disturbance. Due to technical issues, sample sizes for videos were n = 20 heat and 17 control nests. We scored the number of nestlings visible on camera, plus each instance of panting, head-out-box-hole, nestling begging and parental provisioning. For all behaviors, we scored the number of nestlings performing the behavior (Woodruff et al. 2023). *Panting* was defined as a >3 sec period of silent mouth gaping, paired with expanding and contracting body movement. Panting was distinguished from begging via (i) these body movements, which are uncommon during begging, and (ii) audio because panting is silent while begging includes vocalization (elaborated in *SI§B). Head-out-box-hole* was defined as a nestling’s body along or touching the front box wall, with its head directed towards the entrance hole and its neck extended level to, or out of the hole. This behavior can co-occur with panting, and it is a measure of space use in which nestlings move closer to cooler ambient air near the entrance hole. Panting and head-out-box-hole data were binned into five-second intervals (max possible = 720 intervals) and scored ‘present/absent’ within each interval. We calculated the mean *begging intensity* of a brood per feed and the *proportion of nestlings that begged* per feed. Per-feed values were then averaged across the duration of the observation. We quantified the total *number of parental feeds* to rule out potential indirect effects on parents. We re-scored a subset of videos (n = 8) to test for scoring reliability. Intraclass correlation coefficients (ICCs) indicated moderate to high repeatability for focal behaviors (0.72-0.97)(Koo & Li 2016). Details in *SI§B* and *Table S1*.

### Nestling sampling

At the end of the heat challenge, we returned to the nest and confirmed all nestlings survived. Then, we banded nestlings with a numbered USGS aluminum band, measured body mass using a digital Ohaus scale (nearest 0.1 g) and flattened wing length using a stopped wing ruler (nearest 0.5 mm). We sampled blood from the largest, median, and smallest nestlings per nest to facilitate analyses on potential mass-related correlates of HSP gene expression, which is known to elevate within 4 hours of heat (Tu et al. 2016; Xie et al. 2014). Nestlings were bled from the alar vein (∼50 uL; latency from nest disturbance to blood on ice: 7:11 ± 0:15 min), except for the median mass individual, which was euthanized via an overdose of isoflurane followed by rapid decapitation and bled from the trunk (latency to euthanasia: 3:17 ± 0:10 min). From euthanized nestlings, we collected additional tissues, including pectoral muscle and brain, which also express HSPs (Woodruff et al. 2022). Samples were frozen on dry ice in the field and stored at -80 °C. Later, we micro-dissected brains into functional regions (Soma et al. 2003), focusing our analyses on the hippocampus (HPC), a brain area that mediates navigation and spatial memory (Bingman et al. 2003; Pravosudov et al. 2006).

### Quantitative PCR

We quantified relative gene expression using RNA extracted from blood (n = 72 heat, n = 62 control nestlings), pectoral muscle (n = 25 heat, n = 20 control nestlings), and hippocampus (n = 25 heat, n = 17 control nestlings). Sample sizes for some analyses are lower than the total number of nestlings bled or euthanized due to insufficient RNA yield. We extracted RNA using Trizol and converted RNA to cDNA using Superscript III (details in *SI§C*). cDNA was run in triplicate in quantitative real-time PCR (qPCR) to measure mRNA abundance of HSP90AA1, which has been robustly linked to heat tolerance (Tu et al. 2016; Xie et al. 2014). We calculated mRNA abundance with the comparative Ct method (2^-Δct^): fold change in expression for the gene of interest normalized to an internal reference gene, MRPS25. MRPS25 was stably expressed across treatment groups (<1 Ct difference, on average). Details on qPCR reactions, thermal profiles, and primers are in *SI§C* and *Table S2*. Plates were balanced by treatment and date. Each plate included intra- and inter-plate controls (a cDNA pool derived from tree swallow RNA). Inter-plate coefficient of variation (CV) was 2.27%, intra-plate CV was 0.55 ± 0.17%. We found no significant treatment effect on HSP90AA1 gene expression in the pectoral muscle or hippocampus, so we do not discuss these tissues further; see *SI§C*. We also found no significant effect of sex among the terminally collected samples (*Table S4*), so we did not pursue this further.

### RNA-seq and differential gene expression

To shed light on heat-sensitive biological processes beyond HSP90AA1, we submitted total RNA from a subset of blood samples for RNA-sequencing at Indiana University’s Center for Genomics and Bioinformatics. This subset included n = 3 nestlings per treatment (one per nest), balanced by brood size and date; focal nestlings were the median mass nestling in their brood. We constructed Illumina TruSeq stranded mRNA libraries, and 75-cycle paired-end reads were obtained using an Illumina NextSeq 500. After cleaning, mapping, and filtering as in (Bentz et al. 2019), we entered 10,395 genes into a differential expression analysis using DESeq2 (version 1.36.0) in R/Bioconductor (R version 4.2.0) (Love et al. 2014); elaborated in *SI§D*. We functionally analyzed differentially expressed genes (DEG) using Gene Ontology (GO) analyses in PANTHER (Mi et al. 2019); we used human reference terms because they are orthologous to, and more complete than, avian references.

### Fledging and recruitment

To explore whether sublethal heat affected the likelihood of reaching later life history milestones, we first measured fledging success. We checked nests around D21 and identified remaining (dead) nestlings based on their numbered USGS band. All ‘missing’ nestlings were assumed fledged (McCarty 2001) because all boxes have predator guards and because dead nestlings older than D12 are nearly adult mass, so they are not readily removed by parents (Winkler et al. 2020). Second, we measured recruitment into the breeding population, devoting substantial effort from March to July the following two years to capture and identify breeding birds, including returning nestlings from the experiment. This approach provides a robust estimate of recruitment for this species (Lombardo et al. 2020) because our extensive study population spans 36.4 km – well beyond typical natal dispersal distances for this species (8.38 km for females and 2.44 km for males, Winkler et al. 2005); elaborated in *SI§E*.

### Statistical analysis

We used R (RStudio 2022.07.1 build 554) and JMP (JMP Pro 16.0.0) to conduct three types of analyses: 1) treatment effects on nest temperature, 2) treatment effects on nestling phenotypes and performance, and 3) predictors of variation in HSP gene expression, within and among nests. Data and model residuals were visually inspected for normality. Unless otherwise stated, models assume gaussian distribution. We ensured that model variables were not multicollinear (variable inflation factors < 3, Fox & Weisberg 2018). We report the variance explained by fixed (R^2^ marginal) and both fixed and random effects (R^2^ conditional) where applicable. We also report effect size (beta estimate, β, or eta squared, η^2^, depending on model type) and standard error (SE) for each fixed effect.

#### Effects on temperature

To predict mean nest temperature during the experiment, we fit a linear model with fixed effects of treatment, mean ambient temperature, and brood size. Brood size was included because additional nestlings in a confined space may increase heat emission and reduce conductance (Webb & King 1983).

#### Effects on nestling behavior, HSP gene expression, and morphology

Every model included treatment and mean ambient temperature. We expected the number of nestlings in the nest to affect phenotypes because, for example, morphology is related to brood size and behavioral counts are related to the number of nestlings visible on the video. Thus, for behavior models, we also included the mean number of nestlings visible, and for other traits, we included brood size. Brood size and number of nestlings visible are correlated (Pearson r = 0.78). For models that included multiple data points per nest (i.e., mass, wing length, blood HSP gene expression), we included a random effect of nest ID.

To test for heat effects on the amount of panting, we ran a negative binomial regression on the count of time intervals *without* panting; this inversed interval data achieved a better model fit. To test for heat effects on the amount of head-out-box-hole, we ran a zero-inflated negative binomial regression on the count of time intervals with head-out-box-hole. Both thermoregulatory behavior models used the *glmmTMB* package (Magnusson et al. 2017). To ease interpretation of figures, we converted the number of 5-second intervals into minutes. To test for heat effects on the proportion of nestling begging, we ran a log-linked binomial regression, which is a robust approach for proportion data (Chen et al. 2017). To test for heat effects on morphology, our dependent variables were body mass and wing length. To test for heat effects on HSP gene expression in the blood, mRNA abundance values were Log_2_ transformed to improve normality and model fit.

#### Effects on fledging and recruitment

These tests used log-linked binomial regressions with a random effect of nest. The likelihood to fledge analysis included n = 155 nestlings that were not terminally collected. Because the sample size of nestlings that did not fledge was only 3, our model solely tested the main effect of treatment. The likelihood to recruit analysis included n = 152 nestlings that fledged, 13 of which later recruited as adults. For the recruitment analysis, we tested fixed effects of treatment, mean nest temperature, and the interaction between the two.

#### Predicting variation of HSP gene expression

To begin, we quantified the scope of within- and among-nest variation in blood HSP gene expression, using coefficients of variation (CVs). We then tested potential predictors of heat-induced HSP variation, log2-transformed to meet model assumptions. First, using individualized data for all heat-exposed nestlings, we tested for an effect of nest ID using a simple ANOVA. Next, including a random effect of nest ID, we tested whether D12 mass predicted HSP gene expression, using a linear model (LM). Finally, we explored fixed effects of nest temperature, brood size, amount of panting, and amount of head-out-box hole in separate LMs. Because these were measured at the nest-level, our dependent variable was nest-averaged HSP gene expression.

## Results

Nest temperatures were significantly higher in heated nests by an average of 4.5°C (*Fig. 1, Table 1-2*). Nest temperatures were higher at higher ambient temperatures (*Fig. 1*) but were unrelated to brood size (*Table 2*); model: R^2^m = 0.72.

**Table 2:** Model results. Beta estimate effect sizes (β), standard error (SE), test statistic (either F-ratio or z-value depending on model type), and p-value are reported. P < 0.05 are bolded.

| Variable | Predictor | $\beta$ | SE | Test statistic | p value |
| --- | --- | --- | --- | --- | --- |
| Nest Temp | Treatment | 4.05 | 0.49 | $F_{1,42} = 86.18$ | **< 0.001** |
| | Ambient Temperature | 0.32 | 0.06 | $F_{1,42} = 29.04$ | **< 0.001** |
| | Brood Size | 0.36 | 0.24 | $F_{1,42} = 2.32$ | 0.14 |
| Intervals Without Panting | Treatment | -0.63 | 0.25 | $z = -2.54$ | **0.01** |
| | Ambient Temperature | -0.14 | 0.02 | $z = -5.65$ | **< 0.001** |
| | # of Nestlings Visible | -0.04 | 0.13 | $z = -0.33$ | 0.74 |
| Intervals with Heat-out-box-hole | Treatment | 1.32 | 0.60 | $z = 2.21$ | **0.03** |
| | Ambient Temperature | 0.11 | 0.05 | $z = 2.10$ | **0.04** |
| | # of Nestlings Visible | 0.63 | 0.21 | $z = 3.04$ | **0.002** |
| D12 Mass | Treatment | -1.02 | 0.42 | $F_{1,41.71} = 5.87$ | **0.02** |
| | Ambient Temperature | 0.08 | 0.10 | $F_{1,41.30} = 2.65$ | 0.11 |
| | Brood Size | -0.80 | 0.21 | $F_{1,42.69} = 15.13$ | **< 0.001** |
| D12 Wing Length | Treatment | -1.37 | 1.03 | $F_{1,40.70} = 1.78$ | 0.19 |
| | Ambient Temperature | 0.02 | 0.13 | $F_{1,40.31} = 0.03$ | 0.87 |
| | Brood Size | -0.80 | 0.50 | $F_{1,41.62} = 2.51$ | 0.12 |
| *Table 2 (continued)* |  |  |  |  |  |
| BL HSP Gene Expression | Treatment | 1.32 | 0.32 | $F_{1,41.33} = 17.10$ | **< 0.001** |
| | Ambient Temperature | 0.05 | 0.04 | $F_{1,41.61} = 1.54$ | 0.22 |
| | Brood Size | -0.02 | 0.16 | $F_{1,41.84} = 0.02$ | 0.90 |
| Fledging | Treatment | -0.51 | 1.24 | $z = -0.41$ | 0.68 |
| Recruitment | Treatment | 47.08 | 19.75 | $z = 2.38$ | **0.02** |
| | Mean Nest Temperature | 0.31 | 0.38 | $z = 0.81$ | 0.42 |
| | Treatment*Mean Nest Temperature | -1.17 | 0.51 | $z = -2.29$ | **0.02** |

Amount of panting was significantly higher in heated nests (*Fig. 2A*) and on warmer days, but was unrelated to the number of nestlings visible (*Table 2*); model: R^2^m = 0.56. Amount of head-out-box-hole was significantly higher in heated nests (*Fig. 2B*), at higher ambient temperatures, and when more nestlings were visible (*Table 2*); model: R^2^m = 0.50. There was no effect of heat on begging intensity (β = -0.30, SE = 0.24, F_1,33_ = 0.03, p = 0.86; *Fig. S4A*), the proportion of nestlings begging (β = 0.39, SE = 1.65, z = 0.24, p = 0.81; *Fig. S4B*), or amount of parental provisioning (β = 0.38, SE = 5.10, F_1,33_ = 0.28, p = 0.60; *Fig. S5*); details in *SI§B*.

**Figure 2:**
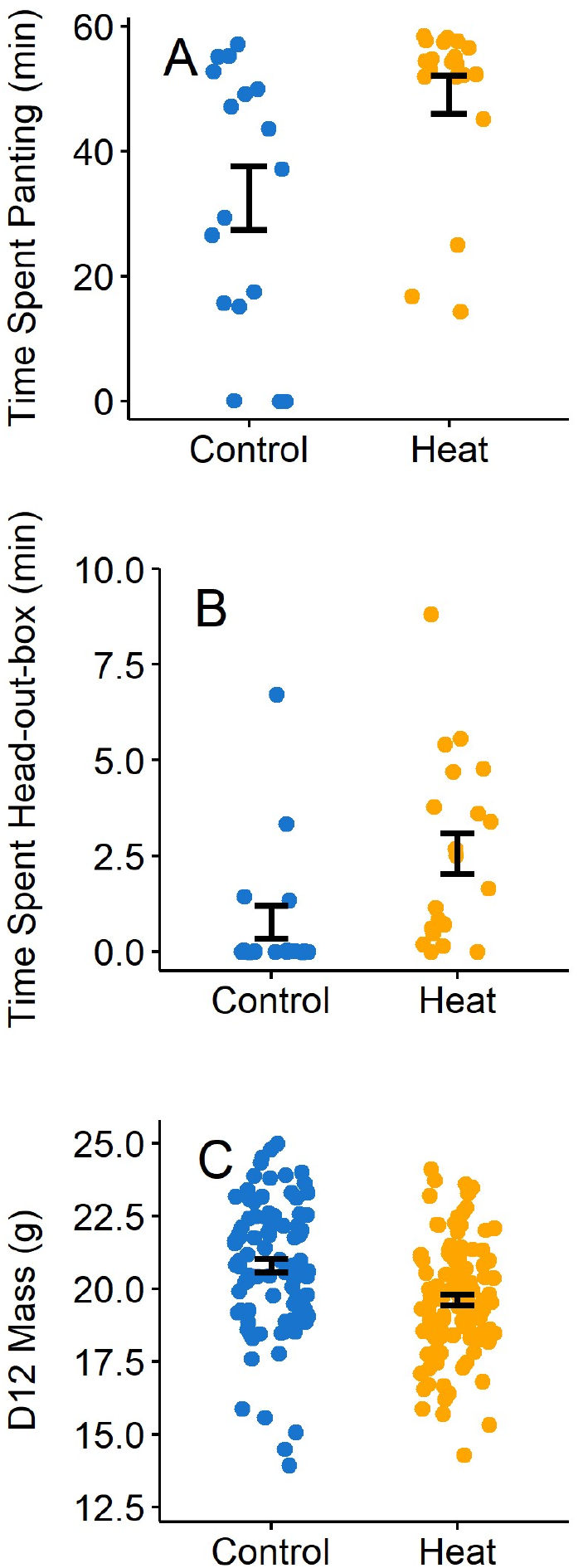
Phenotypic effects of heat. Total duration (minutes) at least one nestling was observed (A) panting or (B) head-out-box-hole during the 1hr observation period. Each point represents one nest. (C) D12 nestling mass (grams) at the end of the experiment. Each point represents one nestling. Mass model accounts for the random effect of nest ID. Error bars are mean ± SE.

Body mass was lower in heated nests (*Fig. 2C*), and in larger broods, but was unrelated to ambient temperature (*Table 2*); model: R^2^m = 0.23, R^2^c = 0.54. On average, heat-exposed nestlings were 19.6 ± 0.2 g and control nestlings were 20.8 ± 0.2 g, a difference of 1.2 g or 5.8%. Wing length was unrelated to treatment (*Fig. S6*), ambient temperature, or brood size (*Table 2*); R^2^m = 0.05, R^2^c = 0.45.

Blood HSP gene expression was higher in heated nests (*Fig. 3, inset*) but was unrelated to ambient temperature or brood size (*Table 2*); model: R^2^m = 0.23, R^2^c = 0.62. In controls, mean within-nest CV was 43.0% (range = 8.0 – 84.1%) and among-nests CV was 79.1%. For the heat treatment, within-nest CV was 59.3% (range = 11.6 – 130.1%) and among-nest CV was 159.4%.

**Figure 3:**
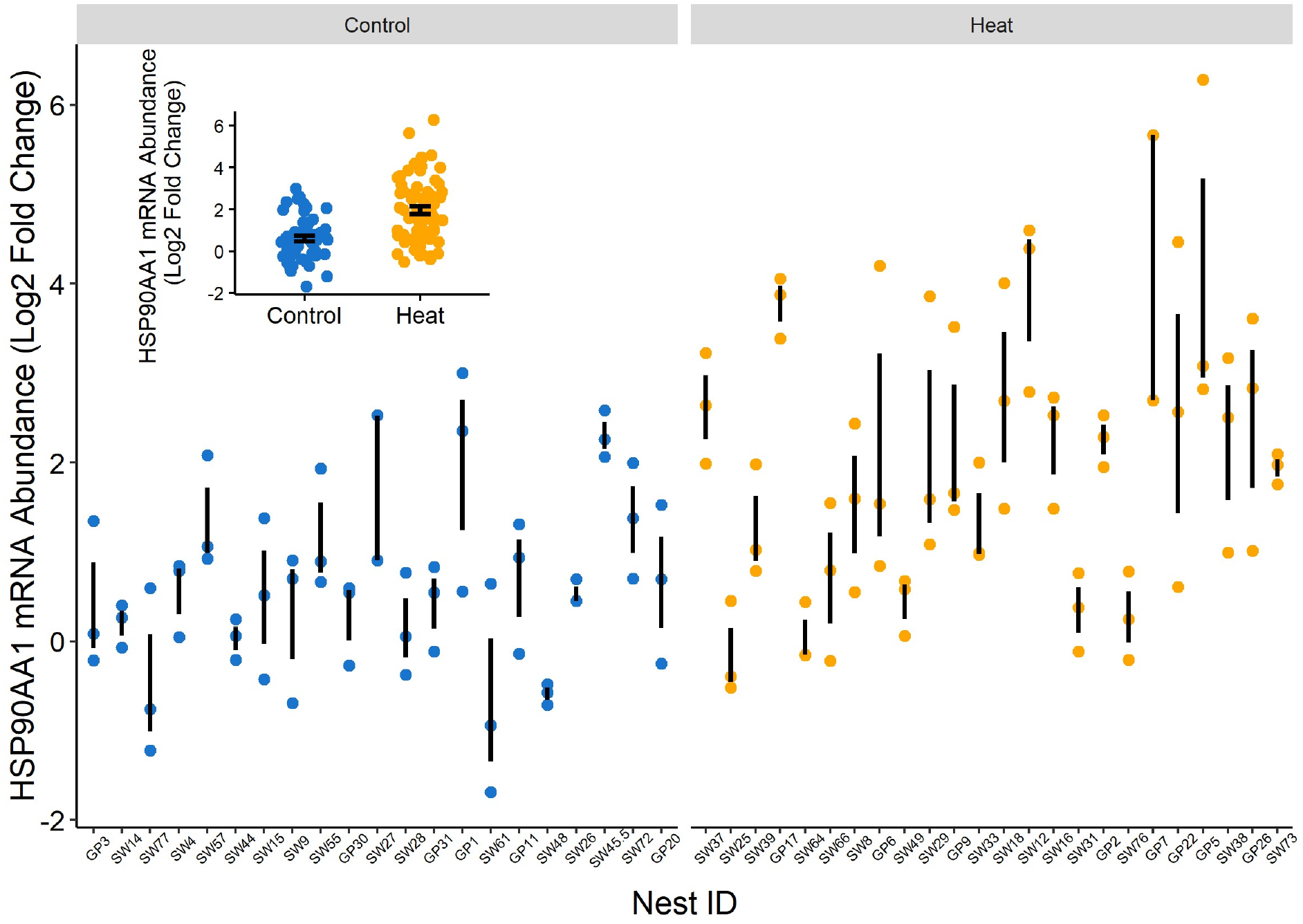
Relative gene expression of blood HSP90AA1 (Log_2_ 2^-Δct^) difference between treatments (inset) and among nests. Within each treatment, nests are ordered by increasing mean nest temperature. Each point represents one nestling. Error bars are mean ± SE for each treatment or nest. Note that 1 unit is a 2-fold difference in abundance on this log2-scale.

Despite this marked variation in heat-induced HSP gene expression, there was a significant effect of nest ID (R^2^m = 0.58, η^2^ = 0.67, SE = 1.06, F_23,47_ = 4.14, p < 0.001). Among heated nests, mean HSP gene expression was unrelated to: nest temperature (R^2^m = 0.08, β = 0.26, SE = 0.18, F_1,23_ = 2.12, p = 0.16), brood size (R^2^m = 0.02, β = -0.17, SE = 0.28, F_1,23_ = 0.38, p = 0.54), panting (R^2^m = 0.01, β = -0.001, SE = 0.002, F_1,18_ = 0.17, p = 0.69), or head-out-box-hole (R^2^m = 0.02, β = -0.01, SE = 0.01, F_1,18_ = 0.32, p = 0.58).

At the individual level, heat-induced HSP gene expression was also unrelated to body mass (R^2^m = 0.02, R^2^c =0.53, β = -0.12, SE = 0.1, F_1,68.43_ = 2.02, p = 0.16); See details in *Table 2* and *Fig. S8*.

We identified 92 DEGs in the blood (0.9% of expressed genes, *Table S5*; see *Table S6* for all 794 DEG prior to false discovery rate correction). HSP90AA1 was the second most affected among these DEG. GO analyses identified several biological processes that were enriched among DEGs, including *protein folding, myeloid cell differentiation, response to heat, response to hormone*, and *negative regulation of apoptotic process* (full list in *Table S7*). Genes within these terms were largely upregulated and included several additional heat shock proteins (e.g., DNAJA1, DNAJB4, HSP90AB1, HSPA2, HSPA4L), plus others related to antioxidants (PRDX4, BIEA, HMOX1, GSTZ1), inflammation (IL1B, TLR2, IFNAR1), metabolism (IRS4, PDK2), and ubiquitination (UBC, MAEA).

Four hours of acute heat had no effect on the likelihood to fledge (*Table 2;* all nestlings fledged except 1 control and 2 heat; R^2^m = 0.02, R^2^c = 0.02). However, recruitment was significantly predicted by the interaction between treatment and nest temperature (*Table 2;* R^2^m = 0.30, R^2^c = 0.30) with higher recruitment among the coolest of the heat-exposed nests (*Fig. S9*). Of the 13 birds that recruited, 10 were from heated nests and 3 were from control nests. This 8.6% recruitment rate is typical for this migratory species (Winkler et al. 2020).

## Discussion

We temporarily elevated nest temperatures by 4.5°C, to an average of 40.9°C, simulating an afternoon we might expect with climate change (Reidmiller et al. 2018). In experimental nests, we documented higher rates of thermoregulatory behaviors but no effect on nestling begging or parental provisioning.

Nestling mass was lower in the heated group despite no treatment differences in wing length, suggesting decreased mass was due to evaporative water loss via panting. Heat also induced high levels of blood HSP gene expression, alongside other transcriptional changes related to antioxidant defenses, inflammation, and apoptosis. HSP90AA1 gene expression was more variable among nests than among siblings in the same nest, and this variation was especially marked after heat exposure. These results shed light on intraspecific variation in the coping mechanisms that may allow for adaptation to rising temperatures.

Behavioral thermoregulation is thought to be a first line of defense against heat (reviewed by: Huey et al. 2012; Muñoz 2022). Heat-induced changes in space use are well documented (Verzuh et al. 2021), but sessile organisms (Pandolfi et al. 2011) and altricial young (Larson et al. 2015) may have limited options. Here, within space-limited nest boxes, heat-exposed nestlings spent more time at the nest box entrance. Nest boxes are naturally hotter than ambient temperatures, so this movement to head-out-box-hole may facilitate access to a cooler microhabitat, in both naturally occurring and experimentally generated heat. Notably, these benefits may not be possible for all nestmates, considering the box entrance is only 4 cm wide, about the same body-width of one nestling. Indeed, across all observations, this behavior was largely limited to just one nestling (∼50% of cases) or two nestlings (∼30% of cases), consistent with observations that access to thermal refuges may be related to social hierarchies (Cunningham et al. 2017). We speculate that, if the benefits of behavioral thermoregulation are density-dependent in confined burrows or nests, then selection may favor smaller brood or litter sizes in a warmer future.

Panting dissipates heat via evaporative cooling and is common across taxa (Loughran et al. 2020; McKechnie et al. 2019). Panting is largely beneficial, dehydration from evaporative water loss represents a real concern. At temperatures experienced during our experiment, water loss in a comparably-sized songbird can be ∼2% of body mass per hour (Albright et al. 2017), though facultative hyperthermia may alter this rate (McKechnie et al. 2019). We observed 1.2g, or 6%, lower mass after ∼4h of heat exposure. We attribute this to evaporative water loss via increased panting because treatments did not differ in structural size (wing length) or apparent food intake (parental provisioning). These results highlight a potential negative effect of heat because large mass is a robust predictor of both lifespan and fecundity (Haywood & Perrins 1992; McCarty 2001); of course, the null effects of heat on begging behavior and fledging success indicate that these effects are short-lived, at least when heat is short-lived.

Damage mitigation and recovery from heat are thought to be regulated by diverse physiological processes, which we documented here. For example, the upregulation of HSP90AA1, as well as other HSPs, should limit cellular damage from heat (Feder et al. 1999; Lindquist et al. 1988). However, sustained heat can outpace the capacity of HSPs to effectively manage denatured proteins. Oxidative damage can trigger the upregulation of genes responsible for the neutralization of free radicals (e.g., PRDX4, BIEA, HMOX1, GSTZ1), and ubiquitination, the marking of damaged elements for destruction (e.g., UBC, MAEA). As physiological stress accumulates, pro-inflammatory responses (e.g., IL1B, TLR2, IFNAR1) recruit macrophages that break down cells (Watanabe et al. 2019). Given that HSPs have been linked with the ubiquitin-proteasome system (Lanneau et al. 2010), antioxidant regulation (Rahman et al. 2022), and inflammatory responses (Zuo et al. 2016), our transcriptomic data may represent a network of co-regulated genes that operate alongside HSPs to combat the deleterious effects of heat.

Re-focusing on HSP gene expression, we documented marked variation within the population, particularly within heat-exposed nests. In terms of fold-differences in gene expression, nestmates differed from one another by an average of 2.6-fold in controls (range: 1.2 to 5.4-fold), but unmanipulated individuals differed by as much as 26-fold across the population. After heat exposure, these numbers grew: heat-exposed nestmates differed from one another by an average of 4.3-fold (range: 1.3 to 14.6-fold), and heat-exposed individuals in the population differed by as much as 112-fold. This high variance after heat exposure is consistent with the idea that environmental stressors may reveal cryptic phenotypic variation (Tanner et al. 2022). Because natural selection can erode trait variation (Wright 1949), it is possible that milder temperatures in the past put more selective pressure on baseline (control) HSP levels compared to heat-induced levels. As heatwaves intensify and become more frequent, this marked variation should be exposed to heat more frequently, though populations with a high degree of standing variation should fare better in the face of climate change (Hoffmann & Sgrò 2011).

We tested several hypotheses on the potential predictors of variation in heat-induced HSP gene expression, noting a significant effect of nest ID. One possible source of this variation is the thermal regime. However, subtle variation in temperature among heated nests did not predict among-nest differences in HSP mRNA abundance. We also did not find a relationship with brood size, even though crowding may increase heat retention (Webb et al. 1983). The degree of behavioral thermoregulation was likewise unrelated to heat-induced HSP gene expression, though we note that the scope of variation in panting was limited due to ceiling effects. Furthermore, we quantified behavior at the nest level (e.g., presence of any panting per time bin), meaning our analyses may not capture among-individual co-variation in behavioral and physiological thermoregulatory mechanisms that has been reported recently (Lipshutz et al. 2022). With these environmental factors accounted for, another possibility is that nest differences in HSP gene expression relate to some genetic component. Thermal tolerance is largely polygenic (Lecheta et al. 2020), and HSP gene expression may have a heritable component (Gleason & Burton 2015). Consistent with this view, we found that nest ID explained 58% of the variation in HSP gene expression; this result provides a maximum possible value for heritability, acknowledging that nestmates may be half-siblings (Whittinghan et al. 2006) and they share much of the same developmental environment, so narrow-sense heritability is surely lower.

Additional findings at the individual level add inferences on sources of variation in thermal tolerance. For example, we expected to see stronger heat responses in larger-bodied individuals because larger bodies retain more heat and can have reduced heat tolerance (Gunderson et al. 2019; Peralta-Maraver & Rezende 2021). However, HSP gene expression was unrelated to body mass, sampling across the full range of size within each nest. Size-dependent effects could be masked if thermal inertia allows larger bodies to heat more slowly (Claunch et al. 2021) or if size determines access to cooler microclimates (Gunderson et al. 2019). Nestmates generally share the same environment, but temperatures can vary across centimeters (Jimenez et al. 2015; Pincebourde et al. 2016). Fine-scale thermal variability within the nest, paired with possible dominance-dependent use of thermal refuges (Cunningham et al. 2017), could facilitate within-nest variation in both the degree of heat challenge and the ensuing responses, though our data did not record at this scale. Our findings nevertheless reject several possible candidates that are hypothesized to drive variation in thermal tolerance, underscoring the need for further study on potential co-variation between HSP gene expression and individual differences in thermoregulatory behavior.

As an added layer of complexity, the tree swallow breeding range has been expanding *south* in the last few decades, into the warm and humid southeastern United States (Shutler et al. 2012; Wright et al.2019), counter to most species that are shifting to higher latitudes or altitudes (Chen et al. 2011). The ecological drivers of this expansion are still unclear (Shutler et al. 2012; Siefferman et al. 2023), but the pattern suggests that tree swallows may be coping well with some degree of heat. Indeed, previous work has demonstrated that tree swallow reproductive success was not sensitive to heatwaves (Taff & Shipley 2023) and here we found evidence that a single warm afternoon positively affected nestlings’ likelihood to recruit in following years, though recruits were more likely from the coolest heated nests. While the sample size of recruited birds is small, it represents typical recruitment rates for this migratory species (Shutler et al. 2006) and we put substantial effort into monitoring our population (see *SI§E*).

Thus, even though our results implicated that heat-exposed nestlings lost mass while upregulating a suite of physiological and behavioral thermoregulatory mechanisms, they still fared well, up to some limit.

Rising temperatures have the power to drive species declines (McKechnie et al. 2019; Rosenberg et al. 2019), and the scope of individual variation will shape the adaptive potential of species. Our approach, which quantifies behavioral and physiological heat-response mechanisms and assesses potential sources of intra-specific variation, revealed variation in heat-responses upon which selection may act.

## Acknowledgements

We thank S. E. Hengeveld and G. T. Smith for feedback on earlier versions, S. E. Wolf, E. M. George, and A. C. Ritter for support with design and interpretation, and the 2021-2023 Rosvall Lab field teams that monitored our breeding population. We thank the Indiana Department of Natural Resources for access to field sites. The project was funded by grants to MJW from the Indiana Academy of Science and the Amos Butler Audubon Society. Additional support came from NSF (IOS-1942192 to KAR; and DBI-1460949 for the REU support to EAG).

