## Supplemental Information, Methods, Tables for "How altricial birds respond to a heat challenge: organismal perspectives on coping with a future climate scenario": Woodruff et al Altricial Bird Heat Responses SI BioRxiv.pdf

Contents

**Supplemental Information** 2
*A. Nest Environment and Experimental Heating* 2
*B. Behavior* 2
*C. RNA extraction, cDNA synthesis, and quantitative PCR* 3
*D. RNAseq and differential gene expression* 3
*E. Recruitment* 4
**Tables** 5
*Table S1* 5
*Table S2* 5
*Table S3* 5
*Table S4* 5
*Table S5* 6
*Table S6* 6
*Table S7* 6
**Figures** 6
*Figure S1* 6
*Figure S2* 7
*Figure S3* 7
*Figure S4* 8
*Figure S5* 8
*Figure S6* 9
*Figure S7* 9
*Figure S8* 10
*Figure S9* 10
**References** 11

### Supplemental Methods and Information

*A. Nest Environment and Experimental Heating*

To habituate birds to foreign objects, we placed a dummy camera, dummy packs, and iButton temperature loggers in the nest box at least 2 days before the treatment began. To minimize non-experimental variation among nests, we added cellophane to gaps in the nest box that were wider than 2 cm. We also shortened nests that were taller than 5 cm, and added a single layer of poster board paper under nests that were thinner than 2 cm. Nest boxes were otherwise physically similar, built of untreated lumber and using the same design.

Uniheat warmers have been used in previous studies to manipulate nest box temperatures (Albert et al. 2023; Corregidor-Castro & Jones 2021; Dawson et al. 2005; Woodruff et al. 2023), however, most studies use one warmer at a time. Because we were targeting temperatures that simulated a future climate scenario and provided nestlings with a challenge above the approximate upper limit of the thermoneutral zone (~37.5°C, avg for small songbirds, extracted from Appendix S1 in Wolf et al. 2017), we used three heat packs to achieve an intense, but short-term challenge. To confirm effectiveness while ensuring the safety of the birds, we performed pilot testing using empty tree swallow nests that were salvaged from a previous breeding season. These tests mimicked the set-up of the experiment, except that nestlings were not present. These pilot tests showed that three heat packs elevated nest temperatures by 5.1 ± 0.3°C above ambient. Based on comparable nest measurements included in (SI, Woodruff et al. 2023), we expected temperatures in occupied nests to be approximately 34°C. Therefore, a 5°C elevation should be above the approximate thermoneutral zone, but below typical lethal limits (sustained exposure to ≥ 45°C Pollock et al. 2021). With these data, we were confident that we would achieve our goals.

*B. Behavior*

Nestling begging intensity was scored using a postural scale from 0 to 4 (adapted from Pilz et al. 2004; Woodruff et al. 2023). A score of 0 indicated that a nestling did not gape or lift its head or neck in response to a parent – the nestling was non-responsive and, therefore, not begging, but was visible to the observer. A score of 4 indicated that a nestling's body was lifted, mouth gaping, and neck extended. We calculated the mean begging intensity per feed across the brood, and next averaged across all feeds to calculate one mean begging intensity value per video. We calculated the mean proportion of nestlings begging by dividing the count of nestlings begging per feed at a 1-4 intensity by the count of nestlings observed (count of 0-4 begging intensity). We then averaged per-feed values across the 1hr observation.

Intra-rater reliability was determined using intraclass correlation coefficient (ICC) estimates that were calculated using the R package ICC (Wolak 2015). ICC estimates are an index that considers within- and among-sample variation (in this case, within and among nest behavior scores) to provide a value of reliability (Koo & Li, 2016). An ICC estimate > 0.5 indicates moderate reliability, > 0.75 indicates good reliability, and > 0.9 indicates excellent reliability (Koo & Li 2016). *Table S1* details the ICC estimates for each behavior in this study.

Mean begging intensity was unaffected by treatment ($\beta$ = -0.30, SE = 0.24, $F_{1,33}$ = 0.03, p = 0.85) and ambient temperature ($\beta$ = -0.01, SE = 0.03, $F_{1,33}$ = 0.94, p = 0.34), but was significantly higher when more nestlings were visible ($\beta$ = 2.58, SE = 0.13, $F_{1,33}$ = 6.40, p = 0.02); model: $R^2m$ = 0.17. Similarly, mean proportion of nestlings begging was not affected by treatment ($\beta$ = 0.39, SE = 1.65, z = 0.24, p = 0.81), ambient temperature ($\beta$ = 0.02, SE = 0.18, z = 0.09, p = 0.93), or the number of nestlings visible ($\beta$ = 0.19,

SE = 0.92, z = 0.21, p = 0.84); model: $R^2m = 0.03$. The total number of parental feeds was significantly higher in larger broods (β = 5.94, SE = 2.46, $F_{1,33} = 5.82$, p = 0.02), but was not affected by treatment (β =0.38, SE = 5.09, $F_{1,33} = 0.28$, p = 0.60) or ambient temperature (β = 0.10, SE = 0.61, $F_{1,33} = 0.04$, p = 0.85); model: $R^2m = 0.15$.

*C. RNA extraction, cDNA synthesis, and quantitative PCR*

To extract RNA from hippocampus, muscle, and blood samples, we used the phenol-chloroform-based Trizol method, following the manufacturer's instructions. Blood samples were extracted using Phase Lock Gel tubes (QuantaBio, Massachusetts, USA) to improve yield. We resuspended total RNA in water and used an Epoch Microplate Spectrophotometer (Biotek, Vermont, USA) to analyze RNA quality and quantity.

To make cDNA, we then treated 1 µg RNA with RNaseOUT Recombinant Ribonuclease Inhibitor (Thermo Fisher Scientific, Waltham, Massachusetts, USA) and DNase (Promega, Wisconsin, USA) for reverse-transcription using oligo dT primers and Superscript III (Invitrogen, California, USA).

For qPCR, we added 3 µl of 1:50 diluted cDNA, or 3 µl water for NTCs, and 7µl of mix (1.94 µl water, 0.03 µl forward primer, 0.03 µl reverse primer, and 5 µl SYBR) into each well for a 10 µl total. The thermocycling condition was set to: 10 min at 95°C, 40 cycles of 95°C for 30 s, 60°C for 30 s, and 70°C for 30 s. Lastly, a dissociation phase (95°C for 1 min, 55°C for 30 s, and 95°C for 30 s) confirmed single-product specificity.

For 7 of 333 samples, RNA concentrations were low (< 110 ng/µL), so we equalized the amount of material loaded into the qPCR reaction by modifying our cDNA recipe and qPCR dilution. To do this, we used 300 ng of RNA for reverse transcriptase and later during qPCR, we used a 1:15 rather than 1:50 dilution qPCR to account for the lower concentration. All samples fell within the Ct range of our standard curves. Primers were previously validated in D12 blood serial dilutions (*Table S2*).

Hippocampus HSP gene expression was unaffected by treatment (β = -0.06, SE = 0.17, $F_{1,38} = 0.11$, p = 0.74), ambient temperature (β = -0.03, SE = 0.02, $F_{1,38} = 1.87$, p = 0.18), or brood size (β = 0.09, SE = 0.09, $F_{1,38} = 1.00$, p = 0.33); $R^2m = 0.07$.

Pectoral muscle HSP gene expression was also unaffected by treatment (β = -0.32, SE = 0.31, $F_{1,41} = 0.92$, p = 0.34), ambient temperature (β = 0.03, SE = 0.04, $F_{1,41} = 0.46$, p = 0.18), or brood size (β = 0.01, SE = 0.15, $F_{1,41} = 0.002$, p = 0.96); $R^2m = 0.03$. Based on these results, tissues seem to be differentially responsive to heat, suggesting that some may be more protected from thermal challenges than others.

We note that for our terminally collected samples for which we had a confirmed sex via visual inspection of the gonad(s), we found no effect of sex on HSP gene expression, nor a significant interaction between sex and treatment across the three tissues in this study (*Table S4*). Due to temporal and financial constraints, we therefore did not pursue further analyses.

*D. RNAseq and differential gene expression*

Six blood RNA samples were run in RNA-seq for a pilot analysis of global heat effects on gene expression. This subset included n = 3 nestlings/treatment, 1 median mass nestlings per nest, matched by date and brood size. We constructed cDNA libraries using a TruSeq Stranded messenger RNA kit (Illumina). Sequencing was performed using an Illumina NextSEq 500 Kit with a 75-bp sequencing module to generate paired-end reads. The resulting reads were cleaned using fastp (version 0.20.1) with parameters "-l 17 --detect_adapter_for_pe -g -p" (Chen et al. 2018), mapped to the tree swallow transcriptome (Bentz et al. 2019) using Bowtie2 (version 2.4.4; Langmead & Salzberg 2012), then filtered to only include proper pairs, which were sorted and indexed using Samtools (version 1.16.1; Danecek et al. 2021). Approximately 27 million read pairs per sample were mapped to the entire transcriptome, accounting for ∼93% (range 92-93%) of the total trimmed read pairs. When mapped against a high-quality protein-only subset of the transcriptome, ~18 million read pairs per sample mapped directly to a specific gene with high confidence. We identified 144,119 transcripts that aligned to known avian proteins in the NCBI database using BLAST following Bentz et al. (2019). To identify a set of non-redundant transcripts that represented known genes, the transcripts were filtered to generate a high confidence set (with at least 50% coverage of the full-length protein and at least 70% identity), which yielded 22,825 transcripts. Putatively unspliced introns and largely redundant transcripts were removed, resulting in a set of high confidence transcripts corresponding to protein-coding segments (n=14,717). Of these, 10,395 genes were found abundantly in blood and used for this analysis. Differential expression analysis was performed using the DESeq2 package (version 1.36.0) in R/Bioconductor (R version 4.2.0) (Love et al. 2014). Transcripts with fewer than 5 total reads across all samples were filtered out. $P$ values were corrected using Benjamini–Hochberg corrections, and FDR ≤ 0.05 were considered differentially expressed.

*E. Recruitment*

We dedicated considerable effort in subsequent years (2022 and 2023) to track recruitment by capturing and identifying all the birds breeding in our study population. From March-July, a large personnel team monitored the ~300 nest boxes in our population (~1000 total person-hours per year). We performed weekly nest box checks until a tree swallow was determined to have established a territory (Winkler et al. 2020), at which point we began checking boxes every one to three days. We monitored nest building, egg laying, hatching, and fledging. We used binoculars to identify leg bands on each tree swallow pair. Recruited nestlings can be easily identified, because we only band nestlings with an aluminum ID band on one leg, whereas birds that are captured as adults are banded with both an aluminum band and a color band. Birds that immigrate into our study population are unbanded in early spring then later banded with an ID and color band. We make it a specific goal to capture, identify, and color band returning nestlings every year.

Previous work using these methods suggests that nearly all living birds will be identified (Lombardo et al. 2020; Shutler et al. 2006), unless there are unmonitored sites within the likely dispersal zone. Our research lab monitored ~300 nest boxes per year, spanning 10 sites that are separated by 0.6-27.2 km and cover 36.4 km end to end. These represent the vast majority of nesting cavities in suitable habitat within the typical natal dispersal radius – 8.4 km for females and 2.4 km for males (Winkler et al. 2005). We often captured recruited females at sites adjacent to their natal site and recruited males at their natal site.

The recruitment rates we observed are typical for wild tree swallows (Butler 1988; Lombardo et al. 2020; Shutler et al. 2006; Wolf et al. 2022). Strenuous migration (Lombardo et al. 2020), inclement weather (Shipley et al. 2020), and poor food availability (Berzins et al. 2021; Lombardo et al. 2020), can contribute to mortality and low recruitment rates. Nestling tree swallow recapture rates range from 0.8-12% (Butler 1988), though yearlings have the lowest return rate (Robertson & Rendell 2001). Previous studies have observed 4.7% (Shutler et al. 2006) and 6% (Lombardo et al. 2020) recruitment. In our own breeding population, recruitment is 8.2% when looking across our population over the past 5 years (155 of 1870 fledglings were recaptured as adults).

**Tables**

| Behavior | ICC value | 95% CI |
|---|---|---|
| Number of time intervals with panting | 0.97 | 0.88 - 0.99 |
| Number of time intervals with head-out-box-hole | 0.87 | 0.51 - 0.97 |
| Total number of parental feeds | 0.91 | 0.64 - 0.98 |
| Mean proportion of nestlings beg per feed | 0.72 | 0.16-0.94 |
| Mean begging intensity across feeds | 0.82 | 0.37 - 0.96 |

*Table S1: Results of Inter-rater reliability. Intraclass correlation coefficient (ICC) estimates and their 95% confidence intervals (CI) for the behaviors analyzed in this study.*

| Gene Name | Primer Sequences | Efficiency | Citation |
|---|---|---|---|
| HSP90AA1 | FWD: GCTTCCAGAAGATGAGGAAGAG<br>RVS: GCAGCATGGAGAAGTGACTAA | 102% | (Woodruff et al. 2022) |
| MRPS25 | FWD: ATCACATCCAGCAACCTTTGG<br>RVS: CAGGGAACTTGGCCTTCAATC | 103% | (Woodruff et al. 2022) |

*Table S2: Primer sequences, efficiencies, and citations.*

| Nest Environment Summaries | Control | Heat |
|---|---|---|
| Mean nest % RH | 57.59 ± 3.49% | 51.51 ± 2.35% |
| Min nest % RH | 48.87 ± 3.64% | 41.46 ± 2.31% |
| Max nest % RH | 69.61 ± 3.17% | 65.79 ± 2.77% |
| Mean difference between nest and ambient % RH | -3.24 ± 1.49% | -7.78 ± 2.20% |
| Mean ambient % RH | 60.83 ± 4.32% | 59.29 ± 3.67% |

*Table S3: Relative humidity (% RH). Mean nest values come from 10min iButton values, averaged per box across the duration of the experiment (± standard error). Ambient values from hourly NOAA values; per-box means were averaged for each treatment.*

| Tissue | Predictor | Estimate | Standard Error | F $ratio_{DF}$ | p value |
|---|---|---|---|---|---|
| Hippocampus | Treatment | -0.40 | 0.26 | $0.11_{1,38}$ | 0.74 |
| | Sex | -0.31 | 0.27 | $0.08_{1,38}$ | 0.79 |
| | Treatment*Sex | 0.63 | 0.35 | $3.19_{1,38}$ | 0.08 |
| Muscle | Treatment | -0.03 | 0.47 | $0.92_{1,41}$ | 0.34 |
| | Sex | 0.27 | 0.46 | $0.004_{1,41}$ | 0.95 |
| | Treatment*Sex | -0.49 | 0.63 | $0.59_{1,41}$ | 0.45 |
| Blood | Treatment | 1.44 | 0.52 | $16.6_{1,41}$ | 0.0002 |
| | Sex | 0.27 | 0.51 | $0.23_{1,41}$ | 0.64 |
| | Treatment*Sex | -0.19 | 0.70 | $0.038_{1,41}$ | 0.78 |

*Table S4: Linear model results testing for sex differences in log2 HSP90AA1 gene expression . Treatment estimates relative to controls, sex estimates relative to females.*

See Supplemental Tables File

*Table S5: Significantly differentially expressed genes in nestling blood after FDR. Differential expression relative to controls.*

*Table S6: Significantly differentially expressed genes in nestling blood prior to FDR. Differential expression relative to controls.*

*Table S7: GO terms for differentially expressed genes in nestling blood relative to controls. +/- indicates pathways that were positively or negatively enriched.*

**Figures**

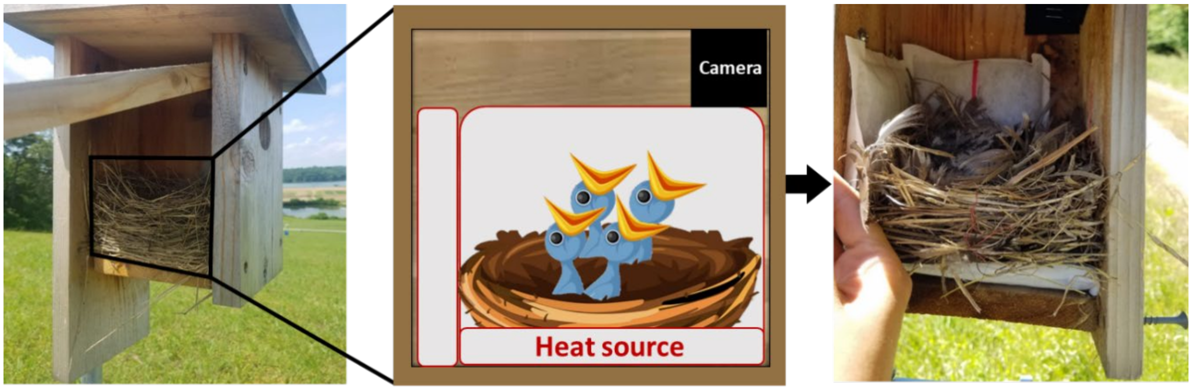


*Figure S1: Experimental set-up.*

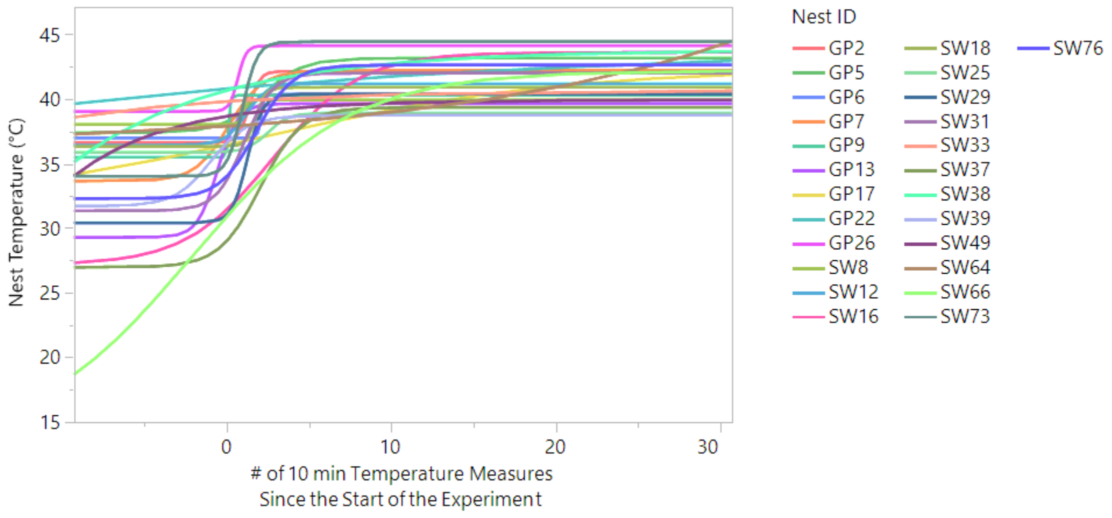


*Figure S2: Nest temperatures (°C) measured before and during the experiment, for heat-exposed nests only. Temperature was measured every 10 minutes (Y-axis). X-axis values denote the number of measurements, such that measurements –9 to 0 correspond to the ten measures just before the experiment, and measures 1-28 correspond to measurements recorded during the experiment. Colors are unique to each nest. Global logistic 4 parameter curve equation for heated nests: 34.4 + ((41.2 – 34.4)/(1 + EXP(-1.3 * (# of 10 min Temperature Measures Since the Start of the Experiment – 1.2)))).*

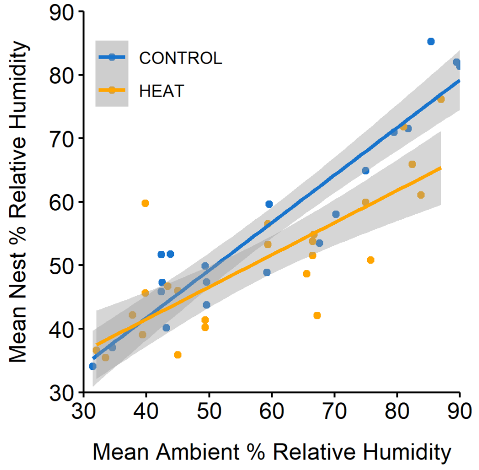


*Figure S3**:** Nest vs ambient percent relative humidity. Data were averaged across the duration of the experiment by treatment groups. Shading is 95% CI.*

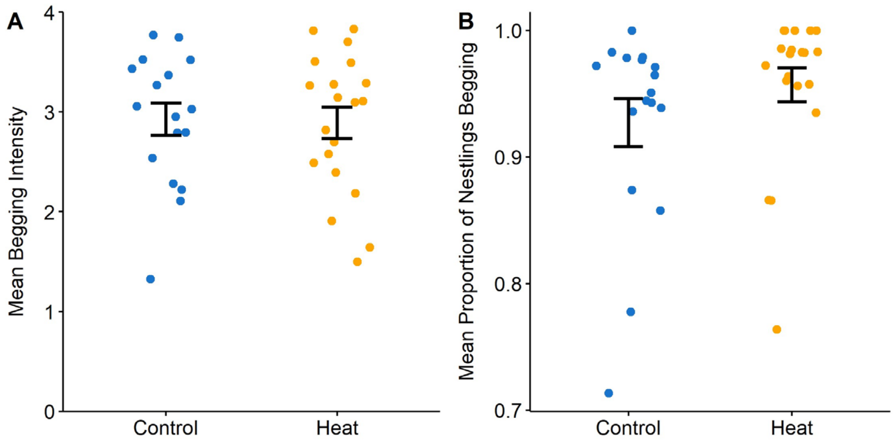


*Figure S4: Treatments did not differ in (A) mean begging intensity (scale 0-4) and (B) mean proportion of nestlings begging per feed during the 1hr observation period. Each point represents one nest. Error bars are mean ± SE.*

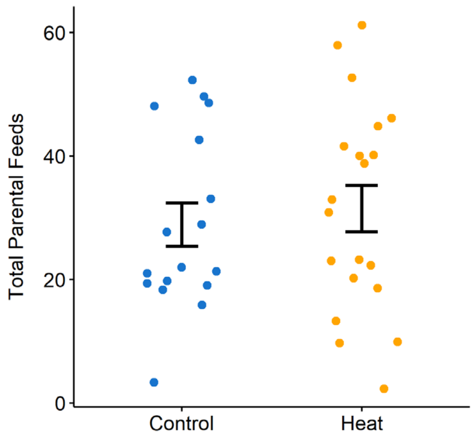


*Figure S5: Treatments did not differ in the total number of parental feeds during the 1hr observation period. Each point represents one nest. Error bars are mean ± SE.*

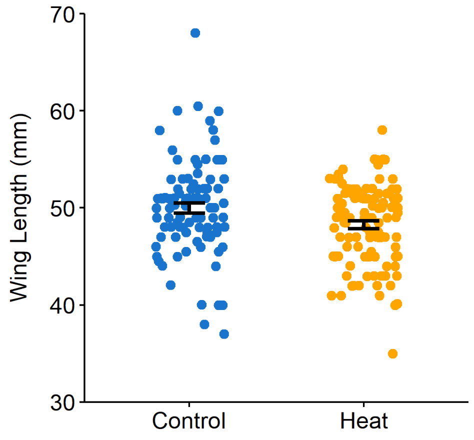


*Figure S6: D12 nestling wing length (mm) at the end of the experiment did not differ by treatment. Model accounts for the random effect of nest ID. Each point represents one nestling. Error bars are mean ± SE.*

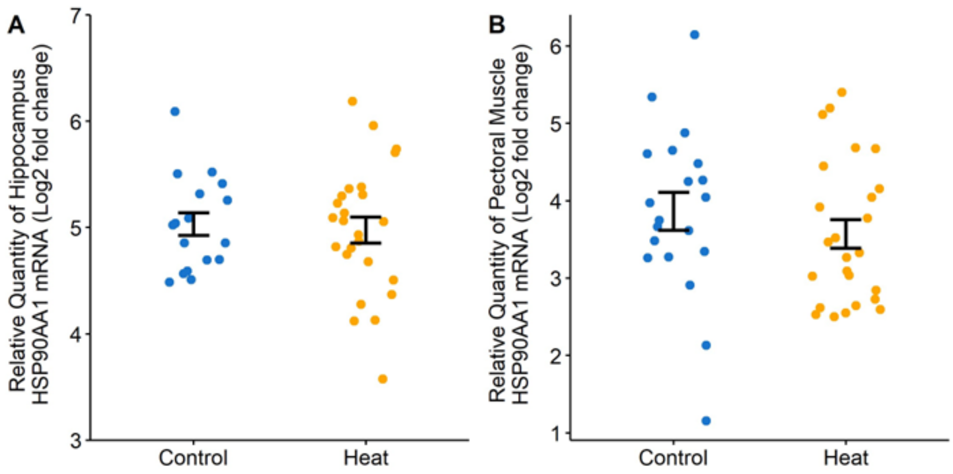


*Figure S7: Treatments did not differ in HSP90AA1 gene expression ($Log_2 2^{-\Delta ct}$), in the (A) hippocampus or (B) pectoral muscle. Each point represents one nestling. Error bars are mean ± SE.*

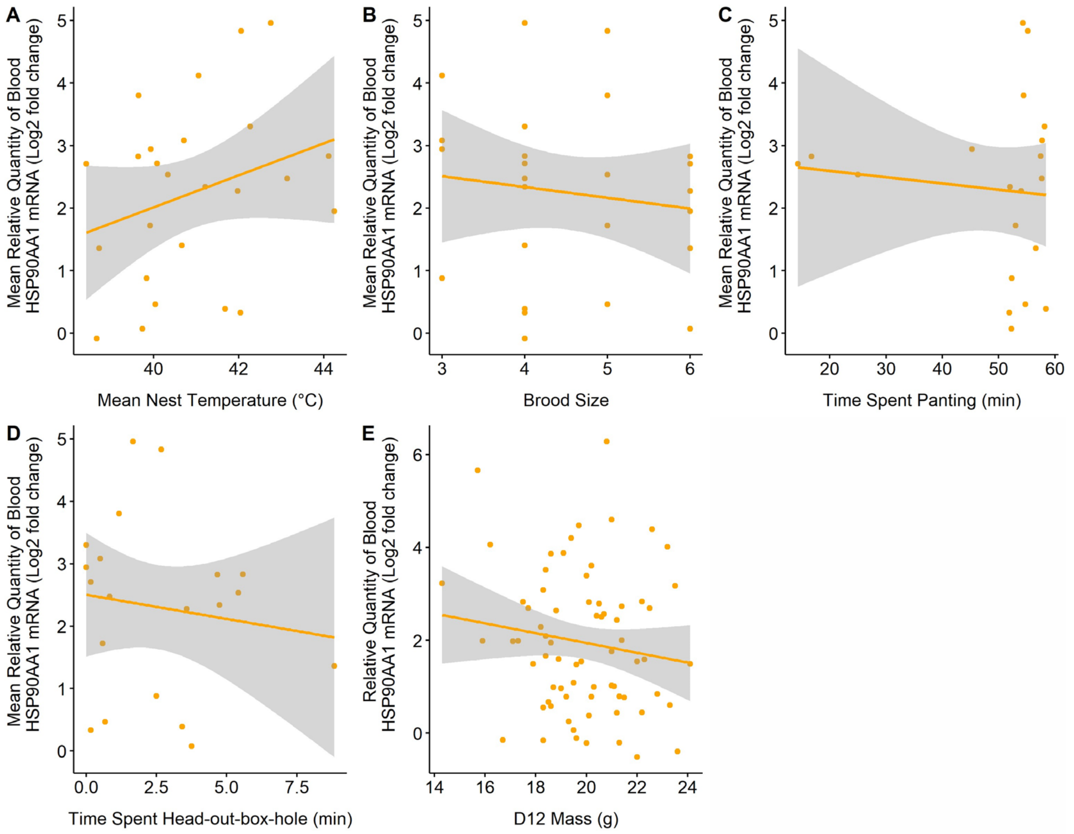


*Figure S8: Heat treatment group blood HSP90AA1 relative gene expression (Log*$_2$ *2*$^{-\Delta ct}$*) by (A) mean nest temperature, (B) brood size, (C) time spent panting, (D), time spent head-out-box-hole, and (E) D12 body mass. Nest-level factors (A-D) are plotted againt mean HSP gene expression. Each point represents one nest. The individual-level factor (E) is plotted against individual HSP gene expression. Each point represents one nestling. Model accounts for the random effect of nest.*

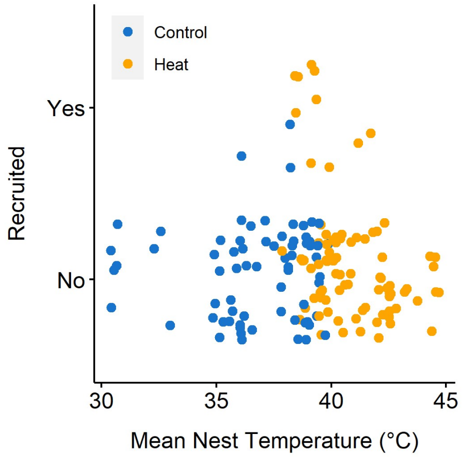


*Figure S9: Relationship nest temperature during the experiment and later recruitment. Each point represents one nestling, and the mean nest temperature in their nest during the experiment.*
